# Lipid membrane topographies are regulators for the spatial distribution of liquid protein condensates

**DOI:** 10.1101/2024.02.20.580889

**Authors:** Chae Yeon Kang, Yoohyun Chang, Katja Zieske

## Abstract

Liquid protein condensates play important roles in orchestrating subcellular organization and in serving as hubs for biochemical reactions. Recent studies have established associations between lipid membranes and proteins capable of forming liquid condensates, and shown that liquid protein condensates can remodel lipid membranes. However, little is known about how the topography of membranes affects liquid condensates. Here, we devised a cell-free system to reconstitute liquid condensates on lipid membranes with microstructured topographies and demonstrated an important role of lipid membranes topography as a biophysical regulator. By employing membrane surfaces designed with microwells, we found that liquid condensates assemble into orderly patterns. Furthermore, we demonstrated that membrane topographies influence the shape of liquid condensates. Finally, we showed that capillary forces, mediated by membrane topographies, led to the directed fusion of liquid condensates. Our results demonstrate that membrane topography is a potent biophysical regulator for the localization and shape of mesoscale liquid protein condensates.

## TEXT

The precise spatial and temporal organization of cellular components plays a fundamental role in numerous life processes, including cell division, cellular migration, and cellular polarization. Traditionally, the plasma membrane and membrane-bound organelles have been considered central hubs of spatial cellular organization. However, in the last decade, a paradigm shift has emerged in our understanding of cellular organization with the investigation of membrane-less organelles, commonly referred to as ‘liquid protein condensates’ or ‘biomolecular condensates’ (1–7). Liquid protein condensates are formed through the assembly of proteins or nucleic acids through unstructured domains or weak multivalent interactions (8, 9).

Both liquid condensates and lipid membranes serve as orchestrators of intracellular spatial organization, and their physical properties and biochemical functions have been the subject of intensive investigation (10, 11). Emerging evidence shows an interplay between lipid membranes and liquid protein condensates (12, 13). For instance, membrane components, such as transmembrane receptors, that are involved in cellular signaling processes are often organized in nano- to micrometer-scale clusters (12). Despite recent progress in understanding these interactions, questions remain regarding the systems-level consequences arising from the interaction between liquid protein condensates and lipid membranes. Specifically, the question of how geometric features of lipid membranes affect liquid protein condensates is still understudied (14). Recent studies describe the remodeling of lipid membranes by liquid protein condensates. Examples of such findings include the observation of liquid protein condensates remodeling plant vacuolar membranes (15) and the remodeling of membranes by endocytic protein condensates (16). In addition, previous studies demonstrated a role of lipid membranes in modulating the concentration threshold for condensate formation and controlling the size of liquid protein condensates. This modulation occurs through the enrichment of condensate components and by limited diffusion (17).

Biomolecular wetting phenomena are emerging as an additional framework for the organization of liquid protein condensates at biological interfaces (14). Notable examples include the regulation of autophagy (18), the role in forming domains (19), and the association with various diseases (20–24). Thus, the hypothesis arises that membrane topography and associated capillary forces may be regulatory factors in the organization of liquid protein condensates. However, a major factor contributing to the limited exploration of how membrane shape influences condensate assembly and distribution is the inherent complexity of living cells. The large number of molecular interactions that occur within cells and the complexity of cellular shapes represent a challenge in systematically unraveling the role of membrane topography for liquid protein condensate regulation.

Cell-free systems are intriguing tools that offer the advantage of disentangling the complexity of living cells, allowing a systems-level exploration of the biophysical mechanisms underlying molecular organization. Consequently, the cell-free reconstitution of biological components in precisely controlled environments is a promising strategy for the systematic investigation of protein interactions with lipid membranes. Recent cell-free studies probed the interaction of liquid protein condensates with supported lipid membranes (9) and giant unilamellar vesicles (25, 26) and revealed, for instance, that liquid protein condensates on supported lipid membranes have the capacity to promote local assembly of cytoskeletal structures (27, 28). Other cell-free experiments demonstrate the role of liquid protein condensates in bending and remodeling lipid membranes (29, 30). These experiments on spherical vesicles established wetting as an efficient mechanism for membrane deformation. However, spherical and flat membranes, due to their uniform topography, may not fully capture the nuanced effects of cellular topography. A controlled assay for the systematic study of liquid condensate assembly in the context of various lipid membrane topographies is required to study condensates in the context of membranes mimicking cellular topographies.

Here, we explored the intricate assembly dynamics of liquid condensates on topographically structured membranes in a well-controlled environment. We developed a cell-free system comprised of topographically structured, supported lipid membranes, and membrane interacting liquid protein condensates, and demonstrate three main findings. First, we show that liquid protein condensates preferentially assemble at the periphery of microstructured membrane compartments. This preference underscores the role of membrane topography in governing the assembly of liquid protein condensates through capillary forces. Second, we demonstrate that liquid condensates deform within the confines of membrane-clad microgrooves. These observations demonstrate the regulation of condensate shape by membrane topography. Finally, our experiments reveal the presence of directionally defined forces acting upon liquid protein condensates in the context of topographically structured membranes. We demonstrate that liquid protein condensates move towards specific locations, defined by the specific arrangements of membrane topography and liquid condensates. These orchestrated interactions result in the emergence of spatial condensate patterns, offering insight into the underlying mechanisms for generating intracellular order.

### Reconstituting liquid protein condensates on membrane-clad microtopographies

We were intrigued by the question of how variations in cell shape may influence processes associated with liquid condensates. However, despite the demonstration of liquid protein condensates occurring at cellular membranes and suggestions that capillary forces due to membrane topography play a role in their organization (14), an experimental system for the systematic investigation of liquid protein condensates in the context of topographically structured membranes was still missing. To address this gap in studying the organization of liquid protein condensates, we developed a cell-free approach designed to reconstitute liquid protein condensates on supported lipid membranes displaying defined topographical structures. Employing photolithography and soft molding techniques, we engineered a thin polydimethylsiloxane (PDMS) layer on top of a glass cover slip (31), featuring cylindrical microwells. Subsequently, we clad these PDMS microstructures with lipid membranes made of the lipid components DOPC and DGS-NTA. Thereby, DGS-NTA served as an engineering solution for tethering liquid protein condensates to the lipid membrane (Figure 1A - C).

**Figure 1.**
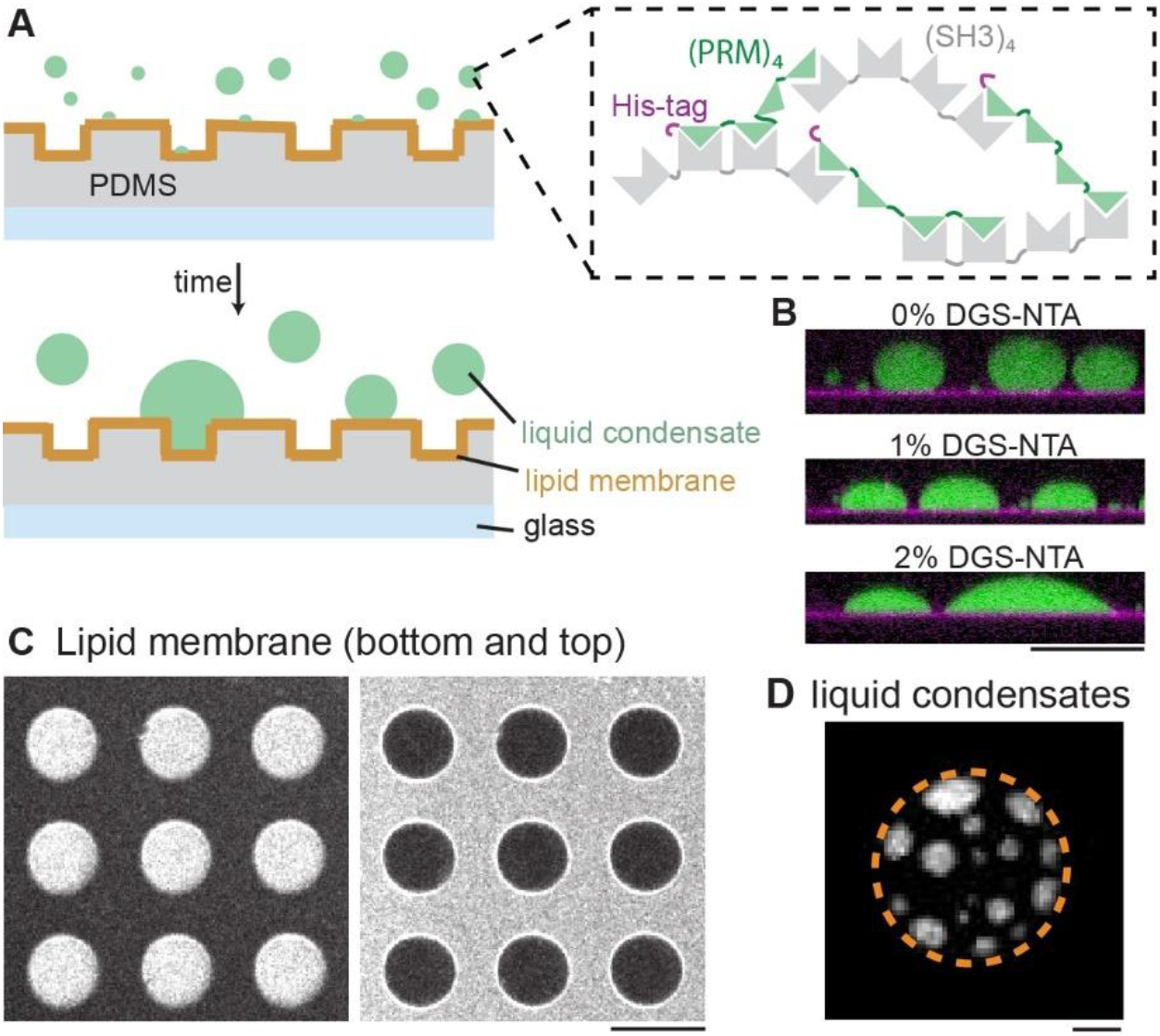
A cell-free assay was established for the spatiotemporal characterization of liquid protein condensates on topolographically structured membranes. (A) Schematic of the experiment. Microstructured PDMS surfaces were clad with supported lipid membranes. The lipid membranes are supplemented with 2% DGS-NTA to mediate the interaction of his-tagged proteins with the membrane. When the proteins PRM4 (50 μM) and SH34 (50 μM) are mixed, they form liquid protein condensates. The liquid protein condensates bind to the membrane through the his-tag of PRM4. (B) Side view of liquid protein condensates (green) on supported lipid membranes with 0%, 1% and 2% DGS-NTA, respectively. Lipid membranes were labeled with 0.05% DiI (magenta). Protein concentrations: 50 μM PRM4, 1% Alexa488-labeled PRM4 (green) and 50 μM SH34. Scale bar: 50 μm (C) Confocal image of a lipid membrane doped with 0.05% DiI, illustrating that lipid membranes are cladding the bottom and top of PDMS microstructures. Scale bar: 50 μm (D) When 50 μM PRM4, 1% Alexa488-labeled PRM4 and 50 μM SH34 were added to microstructured membranes, condensates (white) started to form and attached to the lipid membranes (2% DGS-NTA) through the his-tag of PRM4. The boundary of the microwells are indicated by the dotted orange line. Diameter of microwell: 40μm. Depth of microstructures: 7μm. Scale bar: 10 μm

For the formation of liquid protein condensates, we purified two previously described synthetic proteins (32), each comprising four motif repeats. The first, (SH3)4 has four SRC homology 3 (SH3) domains, while the second protein, PRM4, was composed of four proline-rich motif (PRM) repeats, representing target sequences of SH3 domains. These proteins have previously been demonstrated to form liquid condensates above a critical concentration, and this two-component protein system offers a high degree of control over phase separation behavior through the modulation of protein concentrations and valency (9).

After adding liquid protein condensate components to topographically structured membranes, we confocal microscopy images to verify that liquid protein condensates emerged (Fig. 1D). The emerging protein displayed a characteristic droplet-like appearance and could bind to lipid membranes through the histidine-tag of PRM4. Next, we investigated whether the interaction of condensates and topographically structured membranes affect the distribution and shape of liquid protein condensates.

### Assembly of liquid protein condensates within cylindrical microwells and microgrooves

As liquid condensates exhibit fusion and growth over time, we anticipated the observation of larger condensates over time. To investigate the influence of condensate size on their spatial distribution within cylindrical microwells, we acquired time-lapse confocal microscopy images. As time progressed, we indeed observed a notable increase in condensate volume, together with a reduction in the number liquid protein condensates in individual membrane microwells with a diameter of 40 μm (Fig. 2A, B). The same trend was also evident within smaller membrane microwells of 20 μm (Fig. 2C). Notably, the majority of the liquid protein condensates displayed a peripheral localization pattern after a few minutes. This peripheral localization can be attributed to the larger contact area between liquid protein condensates and lipid membranes at the periphery of the microwells (Fig. 2B) and underscores the impact of membrane topography as a potent biophysical parameter capable of regulating the localization patterns of liquid protein condensates.

**Figure 2.**
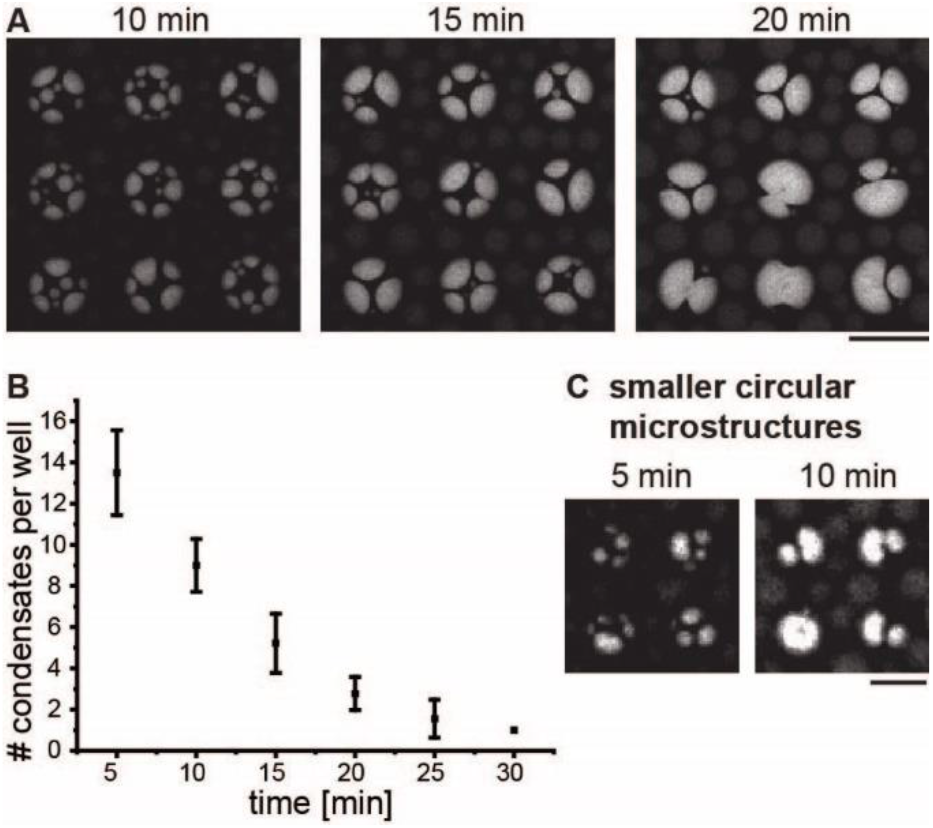
Peripheral assembly of liquid protein condensates within membrane clad microwells. (A) Confocal time-lapse images illustrate the spatial distribution of fluorescently labeled liquid protein condensates within round microwells over time. The liquid protein condensates fuse to generate larger condensates and preferentially localize to the edges of the membrane microwell. Diameter of microwells: 40 μm. Scale bar: 50 μm (B) The number of liquid protein condensates decreases with time until the whole microwell is filled with one large condensate. n>10, error bars: standard deviation. (C) Membrane wells with a smaller diameter of 20 μm are filled earlier than the larger membrane microwell. Depth of microwells: 7μm. Scale bar: 20 μm

Next, to dissect the influence of groove-like membrane topographies, we engineered membrane-clad microgrooves as a topographical feature. We compared the shape of condensates within these membrane grooves in the presence and absence of DGS-NTA. The stretching of condensates was observed on membranes with 2% DGS, but not on membranes without DGS (Fig 3A), demonstrating that the interaction of liquid condensates with lipid membranes plays an important role for the observed shape remodeling of liquid condensates.

**Figure 3.**
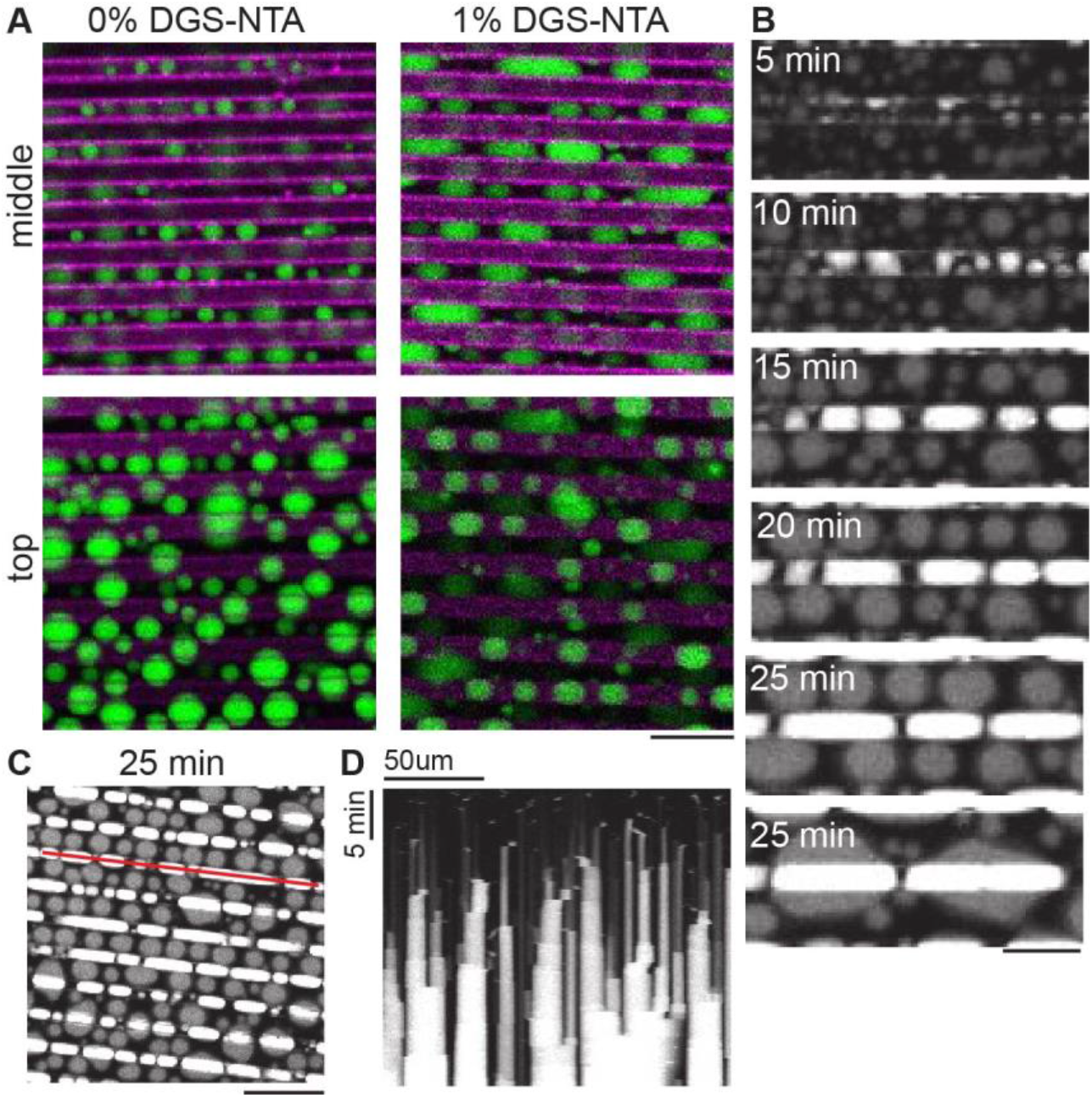
Membrane grooves mediate the stretching of liquid protein condensates along the length axis of grooves in the presence of membrane linkers. (A) Liquid protein condensates (green) were incubated for 15 minutes on grooved membrane surfaces (magenta) containing 0% and 1% DGS-NTA, respectively. Confocal images were acquired on the upper membrane level (top) and 2.5 μm lower (middle). The grooves have a width and depth of 5 μm. The distance between grooves is 5 μm. Scale bar: 20μm (B) Over time, liquid protein condensates outside the grooves and inside the grooves fuse into larger liquid protein condensates. The membrane contact area of lipid condensates outside the grooves remains approximately spherical (grey), the lipid condensates within the grooves (white) are stretched (timepoints: 15 min to 25min). The membrane contained 1% DGS-NTA. Scale bar: 20μm (C) A kymograph was generated at the location indicated by the red line. Scale bar 50μm. (D) Kymograph illustrating the fusion of liquid protein condensates along a membrane groove over time. (B-C) The grooves are 5 μm wide and 7 μm deep. The distance between grooves is 15 μm.

We also compared the shape of condensates within membrane grooves relative to condensates residing on flat membrane regions adjacent to the grooves. Time-lapse confocal microscopy enabled us to image the evolution of liquid protein condensates as they grew over time due to fusion events (Fig. 3B-C).

At the initiation of our experiment, when the droplets were small compared to the width of the grooves, we observed a random distribution of the condensates (Fig 3B, 5 min). However, as the condensates progressively enlarged over time, we noted the cladding of membrane grooves, accompanied by the stretching of liquid protein condensates along the length axis of the grooves. In contrast, the membrane contact area of lipid condensates outside the grooves remained approximately spherical (Fig 3C). The observed stretching of liquid protein condensates within membrane grooves in the presence of DGS-NTA can be attributed to capillary effects, revealing a potential mechanism for the shape adaptability and responsiveness of liquid protein condensates to preformed membrane grooves. Intriguingly, this shape adaptability along membrane grooves also raises the possibility that liquid protein condensates could serve as non-spherical templates upon which forces may act. Pushing forces in the context of cellular membrane grooves, play for example a role in expanding lamellipodial membrane protrusions and cell migration.

### Condensate assembly on membrane topographies smaller than the liquid protein condensates

Thus far, we have characterized the distribution of liquid protein condensates within membrane microwells, and the dimensions of the membrane microwells were larger or comparable to the dimensions of the condensates themselves. However, in living cells, lipid membranes also undergo dynamic topographical changes on the nanoscale. Consequently, liquid protein condensates within cells may encounter topographies smaller than their own dimensions, prompting a fundamental question: How do topographies smaller than liquid protein condensates affect the spatial distribution of liquid protein condensates?

To address this question, we determined whether the distribution of large condensates is affected by small membrane topographies. To do this, we imaged liquid protein condensates that had grown larger than the underlying membrane microstructures and extended across two or more adjacent microwells.

Interestingly, we observed a distinctive phenomenon in which micro-topographies function as pins, that shape the contours of liquid protein condensates (Fig 4A, C). These observations not only underscore the importance of small-scale membrane topographies on the distribution and morphology of liquid protein condensates in the context of model membranes but also provide more general insights into the mechanisms for shape modulation of liquid protein condensates and into how protein architectures interact and respond to membrane features in complex membrane environments.

**Figure 4.**
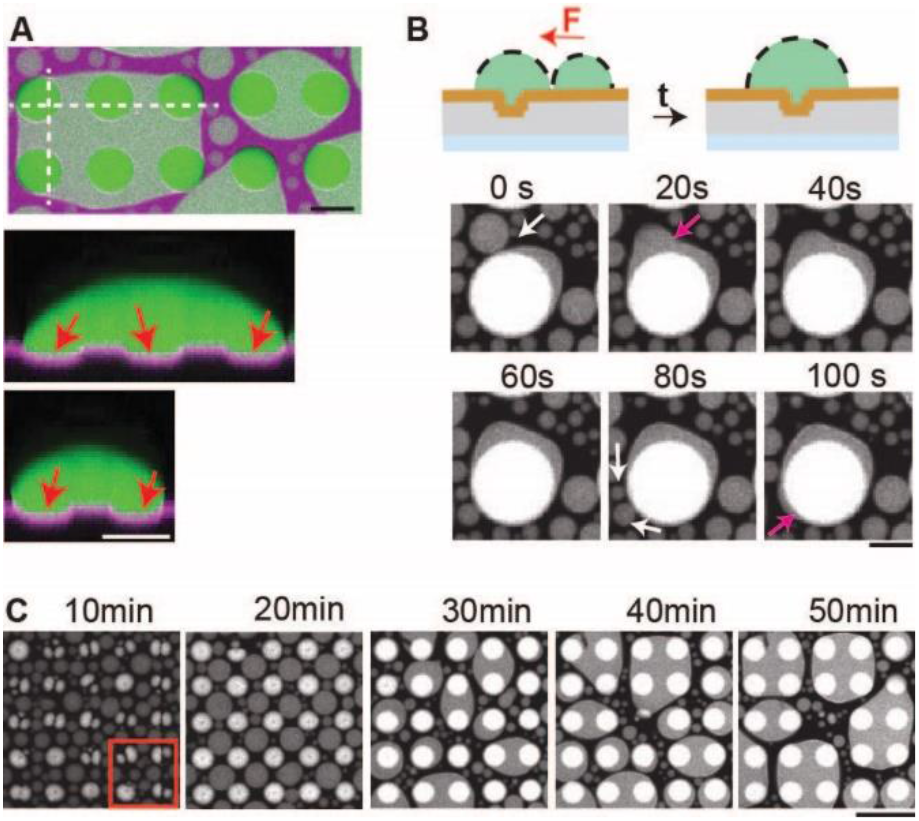
Lipid membrane surfaces with topographical structures modulate larger-scale liquid protein condensates. (A) Liquid protein condensates (green) on top of a microstructured lipid membrane (magenta) and corresponding side views. Diameter of microwell: 20μm. Red arrows point towards microwells. Scale bar: 20μm. (B) Directed fusion of liquid protein condensates towards a liquid condensate, that adheres to a microwell. White arrows: condensate before fusion event. Red arrow: Condensate after fusion event. Diameter of microwell: 40μm. Scale bar: 20μm. (C) Time-lapse confocal images of liquid protein condensates on topographically structured membranes. Red box represents data in figure 2C. Diameter of microwells: 20μm. Depth of microstructures: 7μm. Scale bar: 50μm

In addition to the pinning effect described above, we also observed the phenomena of a directed motion of liquid protein condensates. We attribute this observation to capillary effects within a system of proximal liquid condensates and microwells (Fig. 4B). Forces and capillary effects have previously been postulated as regulatory factors governing the behavior of liquid protein condensates. However, with our assay, we were equipped to experimentally address these phenomena in a controlled way.

## Discussion

In this study, we engineered a cell-free assay to study the interaction of liquid protein condensates with topographically structured membranes by using purified proteins and membrane-clad PDMS microstructures. While biomolecular mechanisms in living cells are often entangled within complex interaction networks of numerous molecular species, our cell-free approach enabled us to study the role of membrane topography for the spatial distribution and shape of liquid protein condensates in a well-controlled environment.

When the protein components PRM4 and SH34 were mixed and added to the membranes, liquid condensates assembled. These condensates attached to the membrane component DGS-NTA through the histidine-tag of PRM4. We studied liquid condensate formation on membranes with user-defined topography of grooves and circular microwells. Using circular microwells, we found that liquid protein compensates preferentially accumulated at the rim of these microstructures. This geometry resembles e.g. the intracellular, bottom contour of adherend cells. Our data suggest that membrane edges may represent effective cues for accumulating or enriching membrane interacting liquid protein condensates. Using grooved membrane topographies, we found that liquid protein condensates stretch along the membrane grooves. The grooved membrane geometries resemble e.g. the intracellular geometry of lamellar membrane protrusions. Our findings suggest, that such cellular membrane geometries may represent effective regulators to mold membrane-interacting liquid protein condensates into non-spherical geometries.

Considering the complexity of living cells, the localization of liquid protein condensates may be determined by additional competing or complementary parameters. For example, in living cells the time and length scales of condensate formation may be specific to condensate composition and other cellular parameters. In other cell-free experiments it has for instance been shown that certain biophysical properties of lipid membranes are regulators for condensate size and assembly: The formation of liquid protein condensates occurs above a concentration threshold and it has e.g. been shown that the recruitment of proteins to lipid membranes can serve as a mechanism for local protein enrichment and subsequent condensate formation at lipid membranes (17). Further, the reduced mobility of membrane-bound proteins, in contrast to freely diffusing proteins, can contribute to the stabilization of relatively small protein condensates (17).

In our experimental system, we employed protein concentrations above the threshold for condensate formation in solution, resulting in relatively large liquid protein condensates. This approach enabled us to investigate the impact of an additional biophysical membrane parameter – namely, membrane topography. The resulting formation of condensates was visualized in 3D on membrane-clad topographies generated by photolithography. Our results display localization patterns of liquid protein condensates in dependence on membrane topography and thus untangle, on a systems level, membrane topography as a general regulator for condensate assembly.

The inherent limitations of minimal model systems, are by definition their inability to mimic the full complexity of a cell and an awareness that minimal systems dissect specific parameters is essential, when working with these systems. In our minimal model system for instance, we reconstituted condensates and engineered microstructures on the order of tens of micrometers. This prompts the question concerning the scalability of our results to the smaller dimensions characterizing cellular condensates and cellular membrane protrusions. The 3D arrangements of condensates within cells often feature dimensions ranging from hundreds of nanometers to a few micrometers. Such small protein condensates are challenging to investigate on membrane topographies using photolithographic methods and standard confocal microscopy, due to the resolution of these techniques. Despite potential non-linear scaling of small liquid protein condensates compared to larger ones, due to different volume-to-surface ratios and surface tension, our model system serves as powerful visualization assay. It enables the experimental demonstration that membrane topography influences condensate localization and offers insights into the trends of how the localization and shape of biomolecular condensates are affected by topographically structured membranes.

In summary, our work bridges concepts of capillary effects with the nanoscale world of lipids and proteins, giving us valuable insights into how molecules are organized by membrane topography – a fundamental biophysical parameter. Our assay represents the first experimental demonstration elucidating localization patterns and shape remodeling behavior of liquid protein condensates on membranes featuring well-defined topographies. They may have further interesting consequences, e.g. for the generation of feedback loops between membrane remodeling processes (26, 33) and condensate remodeling processes. We envision that future miniaturization of membrane topographies, coupled with super resolution imaging and controlled experimental conditions fostering condensate size regulation, hold large promise for unveiling the quantitative dependencies of condensate regulation by membrane topography.

## AUTHOR INFORMATION

### Author Contributions

K.Z. designed research, prepared samples, performed research, analyzed the data and wrote the manuscript. CK supported KZ with sample preparation. YC contributed reagents and acquired microscopy images on non-structured lipid membranes.

### Funding Sources

KZ was supported by the Max-Planck-Society through funding for a Max Planck research group. C.K. was supported by a fellowship (Korea-Germany Science and Technology Fellowship Program.)

### Notes

The author declares no competing interest.

## ACKNOWLEDGMENT

We thank members of the Zieske group for fruitful discussions and Ibrahim el Mazbouh for support during his internship. We acknowledge the Technology Development and Service Group for Nanofabrication’s assistance in providing access to, training in, and management of both the MPL clean room and the nanofabrication technology used for the fabrication of the master template for PDMS molding. The plasmids pMAL-Abl-PRM 4R (Addgene plasmid # 112087) and pGEX SH3(2)-4R (Addgene plasmid # 112090) were gifts from Michael Rosen

## Notes

### Competing Interest Statement

The authors have declared no competing interest.

